# Leveraging AlphaFold2 Structural Space Exploration for Generating Drug Target Structures in Structure-Based Virtual Screening

**DOI:** 10.1101/2025.02.17.638740

**Authors:** Keisuke Uchikawa, Kairi Furui, Masahito Ohue

## Abstract

In early drug discovery, computational virtual screening (VS) is vital for selecting candidate compounds and reducing costs. However, the lack of experimentally determined 3D structures has limited the application of structure-based VS. Advances in protein structure prediction, notably AlphaFold2, have begun to address this gap. Yet, studies indicate that direct use of AlphaFold2-predicted structures often leads to suboptimal VS performance—likely because these structures fail to capture ligand-induced conformational changes (apo-to-holo transitions). To overcome this, we propose an approach that explores and modifies the structural space of AlphaFold2 predictions to generate conformations more amenable to VS. Our method deliberately alters the multiple sequence alignment (MSA) by introducing alanine mutations at key residues in the ligand-binding site, thereby inducing significant conformational shifts. The exploration process is guided by iterative ligand docking simulations, with mutation strategies optimized either by a genetic algorithm or via random search. Our evaluation shows that when sufficient active compounds are available, the genetic algorithm significantly enhances VS accuracy. In contrast, with limited active compound data, a random search strategy proves more effective. Moreover, our approach is particularly promising for targets that yield poor screening results when using experimentally determined structures from the PDB. Overall, these findings underscore the practical utility of modified AlphaFold2-derived structures in VS and expand the potential of computationally predicted protein models in drug discovery.

## 1. Introduction

Drug development and discovery is a costly and time-consuming process. In novel drug development, the estimated cost is several billion US dollars, and it takes more than 10 years to bring a drug to market [1, 2]. One major reason for these high costs is the difficulty of identifying suitable drug candidates. Currently, the total number of drug-like compounds is estimated to range from 10^30^– 10^60^ [3, 4]. However, only a very small fraction of these compounds meet the stringent criteria required for drugs, and many candidates fail during preclinical evaluations [1, 5]. Consequently, selecting promising drug candidates in the early stages of drug discovery is essential to avoid unnecessary expenditures and to promote an efficient development pipeline. Virtual screening, a computational method that identifies hit compounds from a vast chemical space [6, 7], has gained attention. By pre-selecting candidate compounds through virtual screening, unnecessary experimental work can be minimized, potentially leading to substantial reductions in overall development costs and timelines [8]. Virtual screening can be broadly classified into two approaches: ligand-based methods, which rely on known active compounds [9], and structure-based methods, which rely on the protein tertiary structures [8]. In this study, we focused on the structure-based methods.

Structure-based virtual screening (SBVS) is a method of virtual screening that relies on the tertiary structure of a target protein [5, 8]. In SBVS, a compound library is screened by docking ligands into the active site of the protein and ranking them based on their binding affinity calculated using a scoring function. Unlike ligand-based methods that rely on known active compound information, SBVS screens compounds based on the shape of the target protein’s binding site. This approach is considered advantageous for discovering novel drug candidates [5, 10, 11]. It is important to select a suitable protein tertiary structure for SBVS, as its performance is greatly influenced by the structure used. Figure 1 compares screening performance for different CDK2 crystal structures. As shown in Figure 1, even for the same protein, distinct tertiary structures can yield vastly different predictive accuracies. Hence, selecting an appropriate structure for SBVS is of central importance. Generally, it has been reported that the use of a ligand-bound (holo) structure yields higher accuracy than the use of an unbound (apo) structure [12, 13] ^1^ However, the Protein Data Bank (PDB) [14], which is the primary database used in SBVS, does not always contain an appropriate crystal structure of the target protein, particularly the holo structure. Moreover, since experimentally determining the tertiary structure of a novel target protein requires considerable time and cost [15], structural information may sometimes be insufficient.

**Figure 1:**
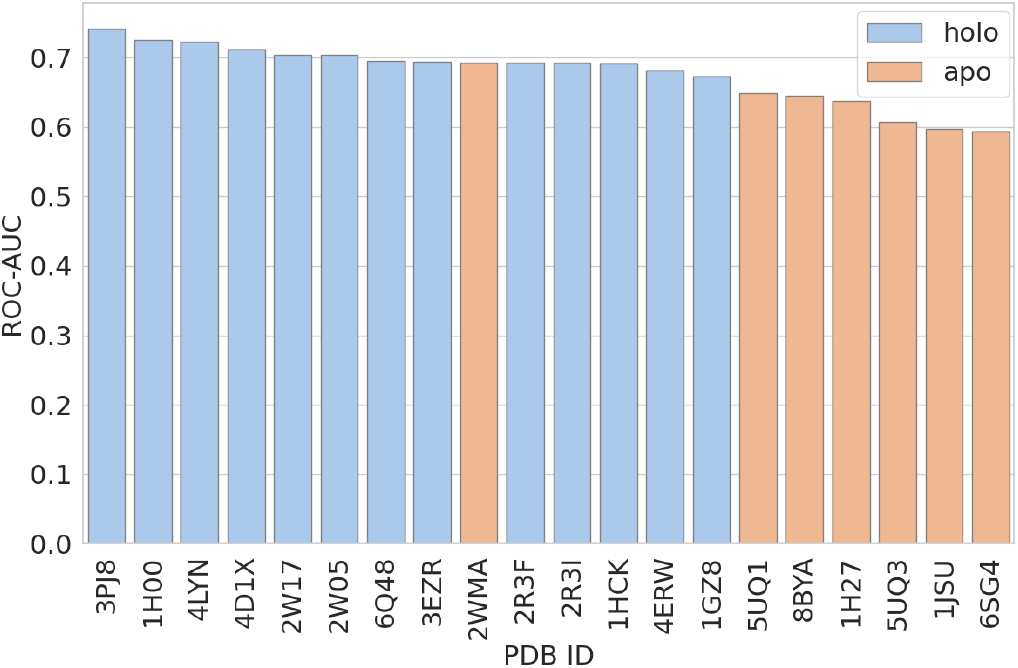
Comparison of screening performance on crystal structures of CDK2 registered in the PDB. The docking simulations using Uni-Dock [16] on the DUD-E dataset [17] were independently conducted us. The performance was evaluated by examining whether the docking score could distinguish active compounds from decoy compounds in the DUD-E dataset.

Under these circumstances, protein tertiary structure prediction technologies have garnered increasing attention. AlphaFold2 [18], a deep learning-based method for predicting protein structures, has achieved groundbreaking results. It predicts the protein tertiary structure directly from its amino acid sequence. One notable feature of AlphaFold2 is its incorporation of co-evolutionary information into a deep neural network model enhanced with attention mechanisms. In particular, by leveraging coevolutionary data extracted from multiple sequence alignments (MSAs), AlphaFold2 achieves highly accurate tertiary structure predictions. The emergence of AlphaFold2 suggests that it may overcome the insufficiency of experimental structural data in SBVS by generating structures that are more suitable for virtual screening.

Although high-accuracy predicted structures are now available thanks to AlphaFold2, previous studies have shown that SBVS using its standard predictions performs similarly to when apo PDB structures are used but worse than when holo PDB structures are used [19, 20]. Therefore, simply using AlphaFold2 predicted structures may not be suitable for SBVS. Various approaches have been explored to refine AlphaFold2 predicted structures for use in SBVS. For instance, Baselious et al. [21] reported enhanced screening performance by refining AlphaFold2 predicted HDAC11 structures through manual adjustments, such as adding ions to the catalytic site, performing energy minimization with known inhibitors, and reorienting side chains in the binding pocket. However, their method lacks general applicability, as it requires target-specific manual modifications. In addition, Zhang et al. [20] improved screening performance by refining AlphaFold2 predicted structures in side chains and loop regions around the binding site, using known holo PDB structures as a reference. However, their method relies on existing holo structural information to guide optimization, which limits its applicability to novel targets since it effectively incorporates partial prior knowledge of the final answer. Therefore, SBVS using AlphaFold2 predicted structures faces several challenges, including the low screening performance with standard predicted structures and the limited general applicability of structure refinement methods. While these approaches focus on refining the predicted structures, there have also been approaches to diversify the outputs generated by AlphaFold2. Typically, AlphaFold2 predicts a single structure from a given input, which limits the range of possible conformations [22]. However, several reports have indicated that adjusting parameters—such as the depth of the MSA or the number of recycling iterations—can introduce diversity into the resulting structures [23, 24, 25]. Still, research aimed at diversifying structures for the purpose of SBVS has not yet progressed sufficiently.

From the current state of SBVS using AlphaFold2 predicted structures, in this study we propose a new approach for generating predicted structures that are suitable for SBVS. Specifically, we vary the parameters of AlphaFold2 to generate diverse predicted structures and perform docking simulations with small-molecule compounds for exploration for each structure. By evaluating the resulting scores, we explore the predicted structures most suitable for SBVS. Furthermore, to evaluate whether the explored structures can achieve generalized high screening performance, we calculate screening performance using them with a separate set of test compounds that were not used in exploration. The objective of this study is to overcome the limitations inherent in the standard predicted structures of AlphaFold2 and to establish a methodology for achieving high-performance and broadly applicable SBVS.

## 2. Materials and Methods

AlphaFold2 leverages information from multiple sequence alignments (MSAs) in its structure predictions, and numerous recent approaches have been proposed that manipulate MSAs to generate diverse structural states [23, 24, 25]. In this study, we focus on the approach by Stein et al. [24], which demonstrated that substituting residues with alanine along the MSA columns can alter inter-residue interactions and thereby produce different predicted structures. This result suggests that this approach may serve as a powerful means for obtaining diverse structure predictions. Building on these findings, we evaluate the impact of various alanine substitution patterns in the MSA on the predicted structures using docking simulation scores with exploration compounds. Based on the results, we explore for alanine substitution patterns that can generate predicted structures suitable for SBVS-structures that enable the accurate evaluation of novel compounds. As a score-based exploration approach, we implemented both random search and a genetic algorithm [26, 27, 28], and evaluated exploration using both a basic search strategy and an optimization strategy. The reason for investigating the genetic algorithm is that the combinations of alanine substitutions in the MSA are enormous, necessitating an efficient method to explore high-scoring configurations. We adopted the genetic algorithm because it can be applied to nonlinear problems with large solution spaces and, since each individual can be evaluated independently, it is well-suited for parallel processing [29]. These characteristics make it appropriate for the present problem, which is nonlinear and computationally intensive. Figure 2 provides an overview of our approach using a genetic algorithm for exploration. First, using AlphaFold2, we generate a set of predicted structures based on MSAs with various alanine substitutions. Next, docking simulations with compounds for exploration are performed on these predicted structures, and each structure is scored. Based on these scoring results, operations such as crossover and mutation are applied to the alanine substitution patterns to generate the MSAs for the next generation. By repeating this process, we aim to generate predicted structures that achieve high screening performance even when evaluated with test compounds that were not used in exploration. In the case of using random search for exploration, the alanine substitution patterns are determined completely at random. In this study, we define “structures suitable for SBVS” as those that can classify active and inactive compounds more accurately in docking simulations.

**Figure 2:**
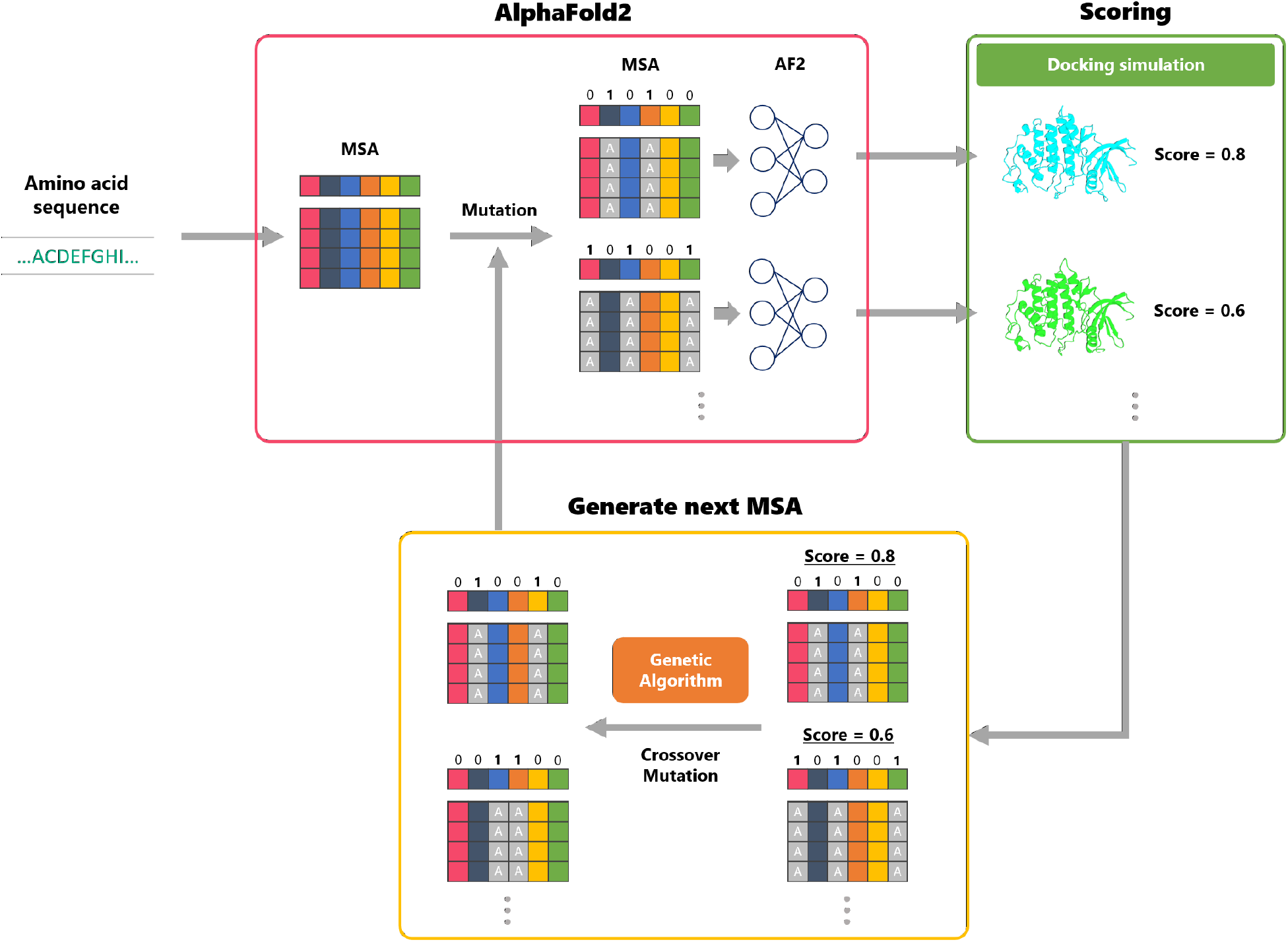
An overview of our approach using a genetic algorithm for exploration

### 2.1. Generating predicted structures with AlphaFold2

We used LocalColabFold [30] to generate predicted structures, as it allows us to specify certain parameters during structure prediction.studies have reported that by reducing both the number of MSAs and the number of recycling steps, one can induce conformational changes in the predicted structure [23]. Accordingly, we fixed the parameters so that LocalColabFold uses shallow MSAs and a minimal number of recycles. The detailed configuration is shown in Supporting Information S1. Input sequences for each target were obtained from UniProt [31]. Furthermore, in using the predicted structures, we trimmed regions with pLDDT *<* 50 and removed parts that were deemed unreliable to prevent disordered regions from interfering with the binding site. The pLDDT score is a residue-level confidence metric output by the AlphaFold model: any residue with pLDDT *<* 50 is considered to have low predictive reliability [18]. If the overall average pLDDT of a predicted structure was below 50, that predicted structure was not used.

### 2.2. Exploration of alanine substitution patterns in MSA

In this study, we explore alanine substitution patterns in the MSA that enable AlphaFold2 to generate structures more suitable for SBVS. Alanine substitution candidates are restricted to residues located within 8 °A of the ligand. This is because such residues are likely to be critical for protein–ligand interactions within the binding site, thus impacting the success or failure of docking. In this experiment, the binding site was treated as known and extracted from the holo PDB structure. Residues selected for alanine substitution were chosen based on the distance between the ligand in the holo PDB structure and the standard AlphaFold2 predicted structure of the UniProt input sequence. If any atom of a residue was within 8 °A of the ligand, the entire residue was considered to be within 8 °A. In our exploration approach, the presence or absence of an alanine substitution is represented as a 1/0 bit in a genetic representation. Concretely, for each column in the MSA generated by AlphaFold2, a bit value of 1 indicates that all residues in that column should be substituted with alanine (2). Our substitution procedure follows the same approach as in the previous study [24]. Alanine was chosen because it primarily disrupts existing interactions while minimizing its overall impact on the protein structure. The detailed configuration is shown in Supporting Information S2.

### 2.3. Scoring method for predicted structures

To perform our exploration approach, we need a mechanism to score how suitable each predicted structure— obtained by mutating the MSA—is for SBVS. In this study, we adopted protein–ligand docking simulations for the scoring process.

#### 2.3.1. Docking simulation settings

For docking simulations, we used Uni-Dock [16], which improves efficiency through GPU usage compared to conventional AutoDock Vina [32], enabling fast and accurate docking. Given the need for docking simulations on a large number of predicted structures, Uni-Dock’s speed is a significant advantage from a computational cost perspective. We defined the docking grid by referring to experimental structure data from the PDB. Specifically, we used PyMOL [33] to extract the ligand’s central coordinates from the holo PDB structure and set the docking grid box accordingly. The grid size was set to 20 °A × 20 °A × 20 °A. Docking with Uni-Dock was performed with the scoring function set to “vina”, the search mode set to “balance”, and the random seed fixed at 1. We superimposed the predicted structure with the holo PDB structure such that the binding site residues within 8 °A of the ligand were aligned, ensuring that the docking center would match between them. For protein and ligand preprocessing, we used AutoDock Tools [34].

#### 2.3.2. Evaluation metrics for predicted structures

We calculated the ROC-AUC value [35] based on the ranking of ligands by their docking scores, taking into account their known active or inactive labels. This ROC-AUC value served as the score for each predicted structure, guiding the genetic algorithm’s operations (e.g., crossover, mutation). The ROC curve is a plot of the true positive rate against the false positive rate, and the AUC is the area under this curve. ROC-AUC values range from 0 to 1, with values closer to 1 indicating higher predictive accuracy. A value of 0.5 or below suggests classification performance equivalent to or worse than random guessing. In the SBVS context, a higher ROC-AUC value indicates that active compounds were more accurately distinguished from inactive compounds by the predicted structure (i.e., that the structure is more suitable for screening). We chose ROC-AUC because it is commonly used for evaluating screening performance and yields values in the interval [0, 1], making it convenient for use as a score in a genetic algorithm.

### 2.4. Dataset

In this study, we selected both proteins that AlphaFold2 has already trained on and those that AlphaFold2 has not trained on as our targets. For targets trained on by AlphaFold2, we used the well-known DUD-E dataset [17], a benchmark dataset for SBVS that includes 102 proteins from diverse classes such as kinases, proteases, nuclear receptors, GPCRs, ion channels, and enzymes. Each target is associated with multiple active and inactive compounds, enabling evaluation of screening performance by measuring binary classification metrics. From this dataset, we selected CXCR4 [36] and KIF11 [37] from the Diverse Subset, as well as the canonical kinase CDK2 [38]. We chose CXCR4 because it yields the lowest screening performance and KIF11 because it yields the highest screening performance within the Diverse Subset [32]. This allowed us to thoroughly test whether our proposed method is effective. CDK2 is a typical kinase with a larger number of known active compounds in DUD-E relative to CXCR4 and KIF11, providing ample data for our experiments. To verify whether our approach can be applied to novel proteins not learned by AlphaFold2, we also selected targets that fall outside of AlphaFold2’s training data. The version of AlphaFold2 (LocalColabFold v1.5.5) used in our experiments was trained on PDB structures deposited up to April 30, 2018 [18]. Therefore, any protein deposited in the PDB after that date can be considered unknown to AlphaFold2. We searched for proteins satisfying the criteria that (1) they were not learned by AlphaFold2 or its close homologs, (2) a holo crystal structure is currently available, and (3) there is sufficient information on active compounds. Consequently, we selected ABHD6 (PDB ID: 7OTS [39]) and HIPK3 (PDB ID: 7O7J) [40] as targets. Their active compounds were obtained from ChEMBL [41, 42], where any compound with an IC_50_ *<* 10 nM was defined as active. We then generated decoy compounds for these active compounds using DUD-E Generate Decoys [17]. The detailed information is shown in Supporting Information S3.

### 2.5. Splitting the ligand data

In this study, the exploration uses an ROC-AUC score based on protein–ligand docking simulations with active and inactive compounds. To examine the effect of the number of known active compounds, we evaluate performance by varying the number of ligands used in the exploration under the following three conditions:

> **dataset-large)** two-thirds of the known active and inactive compounds were used for exploration, while the remaining one-third were reserved for testing.
>
> This setting aims to investigate exploration performance when sufficient ligand data are available. We performed 3-fold cross-validation, splitting the data by clustering based on Tanimoto distances.
>
> **dataset-middle)** 30 known active compounds and 1,500 known inactive compounds were used for exploration, with the remaining compounds reserved for testing. This setting aims to verify exploration performance when available ligand data are relatively limited. We created 3 non-overlapping exploration sets and conducted independent explorations for each set.
>
> **dataset-small)** 10 known active compounds and 500 known inactive compounds were used for exploration, with the remaining compounds reserved for testing. This setting aims to verify exploration performance when available ligand data are limited. We created 3 non-overlapping exploration sets and conducted independent explorations for each set.

For each dataset, we create 3 pairs of exploration and test sets to evaluate generalization screening performance. The exploration and evaluation conducted for each pair are referred to as trial 1, 2, and 3. The 50-fold ratio of inactive compounds to active compounds reflects the design of DUD-E Generate Decoys, which produces about 50 decoys per active compound.

Note that for HIPK3, the number of known active compounds was insufficient, causing the numbers of ligands under the dataset-large and dataset-middle conditions to be nearly the same. Consequently, we use the dataset-large results as a reference for dataset-middle.

### 2.6. Evaluation of the proposed method

To evaluate the performance of our proposed method, we conducted the following experiments.

First, we assessed the screening performance using PDB structures. This allowed us to establish a baseline reference for screening performance against which subsequent results could be compared.

Next, to evaluate the screening performance using AlphaFold2 predicted structures, alanine substitutions were introduced into each column of the MSA with a probability of 0.5, and the average screening performance using the randomly generated structures was calculated. In addition, among the randomly generated predicted structures, we examined the screening performance that achieved the highest results on the test set. Through these two steps, we clarified the representative levels of screening performance using AlphaFold2 predictions can achieve.

We employed random search to explore alanine substitutions in the MSA, aiming to identify parameters that generate predicted structures more suitable for SBVS. As in the previous experiment, alanine substitutions were introduced into each column of the MSA with a probability of 0.5. Subsequently, we performed an exploration with a genetic algorithm to optimize alanine substitutions in the MSA.

Finally, we compared the screening performance of the predicted structures generated by our proposed method (via random search and a genetic algorithm) with that of the existing PDB structures and standard AlphaFold2 predictions. Through this comparison, we comprehensively evaluated the effectiveness of our proposed method and its advantages over alternative approaches.

For all experiments involving predicted structures (excluding those based on PDB structures), we generated and evaluated a total of 1,100 predicted structures to ensure consistency in exploration and assessment.

## 3. Results and Discussion

### 3.1. Comparison of methods based on the achieved screening performance

Figure 3 shows a comparison between the screening performance using the predicted structures explored by the genetic algorithm and random search, and the screening performance using PDB structures and AlphaFold2 predicted structures. First, in all cases, it was confirmed that the predicted structures explored by the proposed method (via random search and a genetic algorithm) were more suitable for SBVS than the average AlphaFold2 predicted structures. Furthermore, for four target (CXCR4, CDK2, ABHD6, and HIPK3) the predicted structures explored by the proposed method tended to be more appropriate for SBVS than the PDB structures. These results demonstrate the effectiveness of the proposed method compared with conventional methods. Notably, the four targets (CXCR4, CDK2, ABHD6, and HIPK3) are those for which the screening performance using PDB structures is relatively low. Therefore, it is considered that the proposed approach effectively enhances screening performance for such targets. Moreover, when comparing Figure 3A, B, and C, it was observed that the screening performance using the explored predicted structures tended to improve as the number of ligands available for exploration increased. In particular, when a larger number of ligands was available for exploration, many cases were observed in which the predicted structures explored by the proposed method achieved test set ROC-AUC values that were comparable to or even exceeded the highest ROC-AUC observed among the AlphaFold2 predictions on the test set. Therefore, the proposed method is considered to be increasingly effective as the number of ligands available for exploration grows.

**Figure 3:**
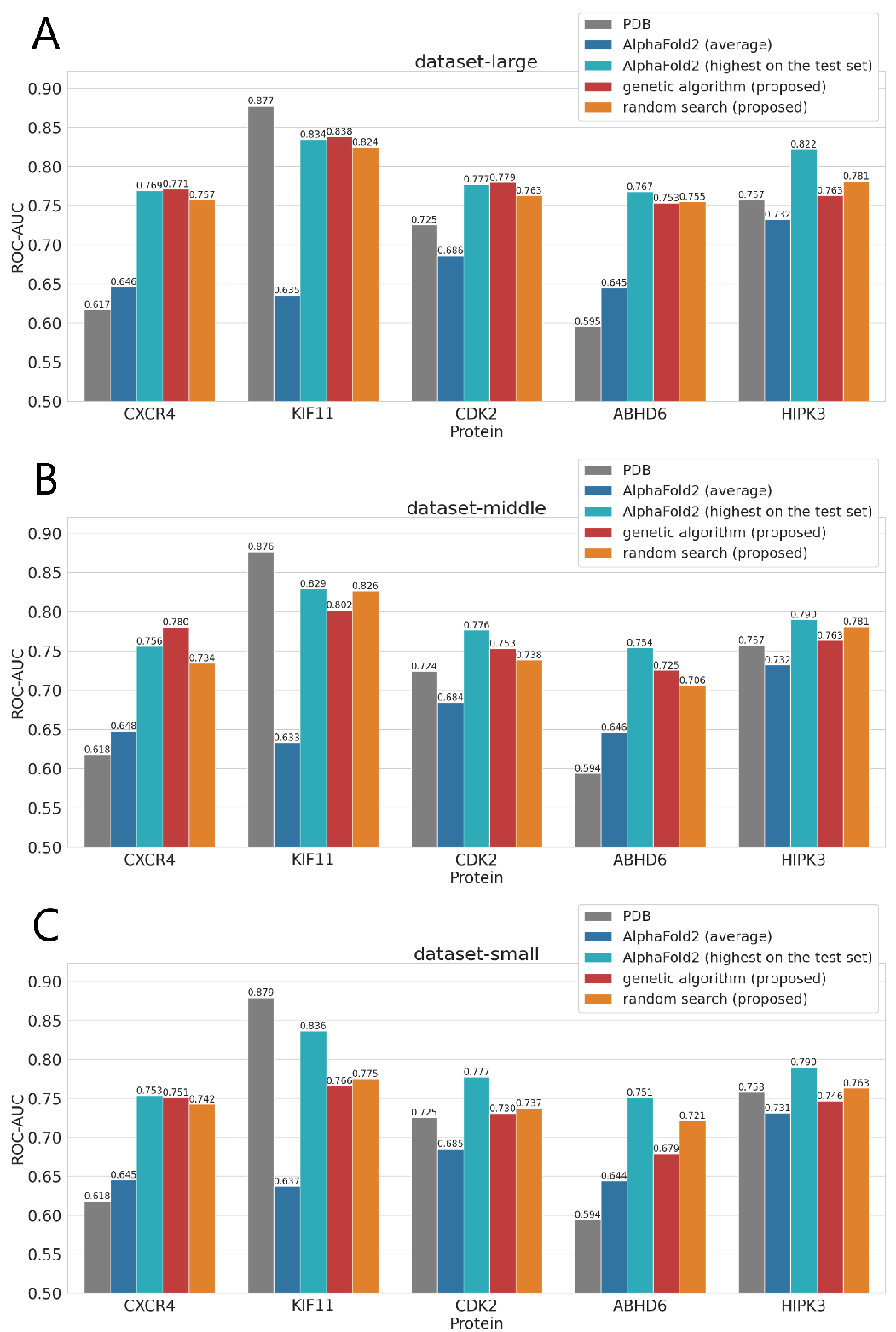
Comparison of screening performance (ROC-AUC) for predicted structures explored by the proposed methods (genetic algorithm and random search) with those of existing PDB structures and standard AlphaFold2 predictions. For all experiments involving predicted structures (excluding those based on PDB structures), 3 trials were conducted, and the average results were used for comparison. A, B, and C show the results for dataset-large, dataset-middle, and dataset-small, respectively.

Comparing the proposed methods, random search and the genetic algorithm, it was observed that the greater the number of ligands available for exploration, the higher the tendency for the screening performance using the predicted structures explored by the genetic algorithm to exceed that achieved by random search. In dataset-large, for three targets with a high number of known active compounds in the exploration set (in descending order: CDK2, KIF11, and CXCR4), the genetic algorithm explored structures that outperformed those explored by random search. These findings suggest that the effectiveness of the genetic algorithm in exploring predicted structures suitable for SBVS increases with the number of known active compounds available. Conversely, when the number of ligands available for exploration is small, random search tended to yield predicted structures with higher screening performance than those explored by the genetic algorithm. In summary, when a sufficient number of known active compounds (30 or more) is available, exploration using the genetic algorithm is effective; however, when only a few active compounds are available, using random search is considered more appropriate.

### 3.2. Transition of exploration in a genetic algorithm

To investigate how the genetic algorithm performed its exploration, we examined the transition of the top ROC-AUC value in each generation, as exemplified in Figure 4. Overall, the results indicate that when the ROC-AUC on the exploration set improved, the ROC-AUC on the test set also tended to improve (Figure 4A). This is considered to be because the greater the number of active compounds available for exploration, the more correlated the ROC-AUC between the exploration set and the test set becomes, thereby enabling appropriate exploration via the genetic algorithm. Furthermore, even when a temporary decline in screening performance is observed, it is thought that the genetic algorithm converges toward a population with a strong tendency for high performance, from which new, more suitable structures for SBVS can be explored.

**Figure 4:**
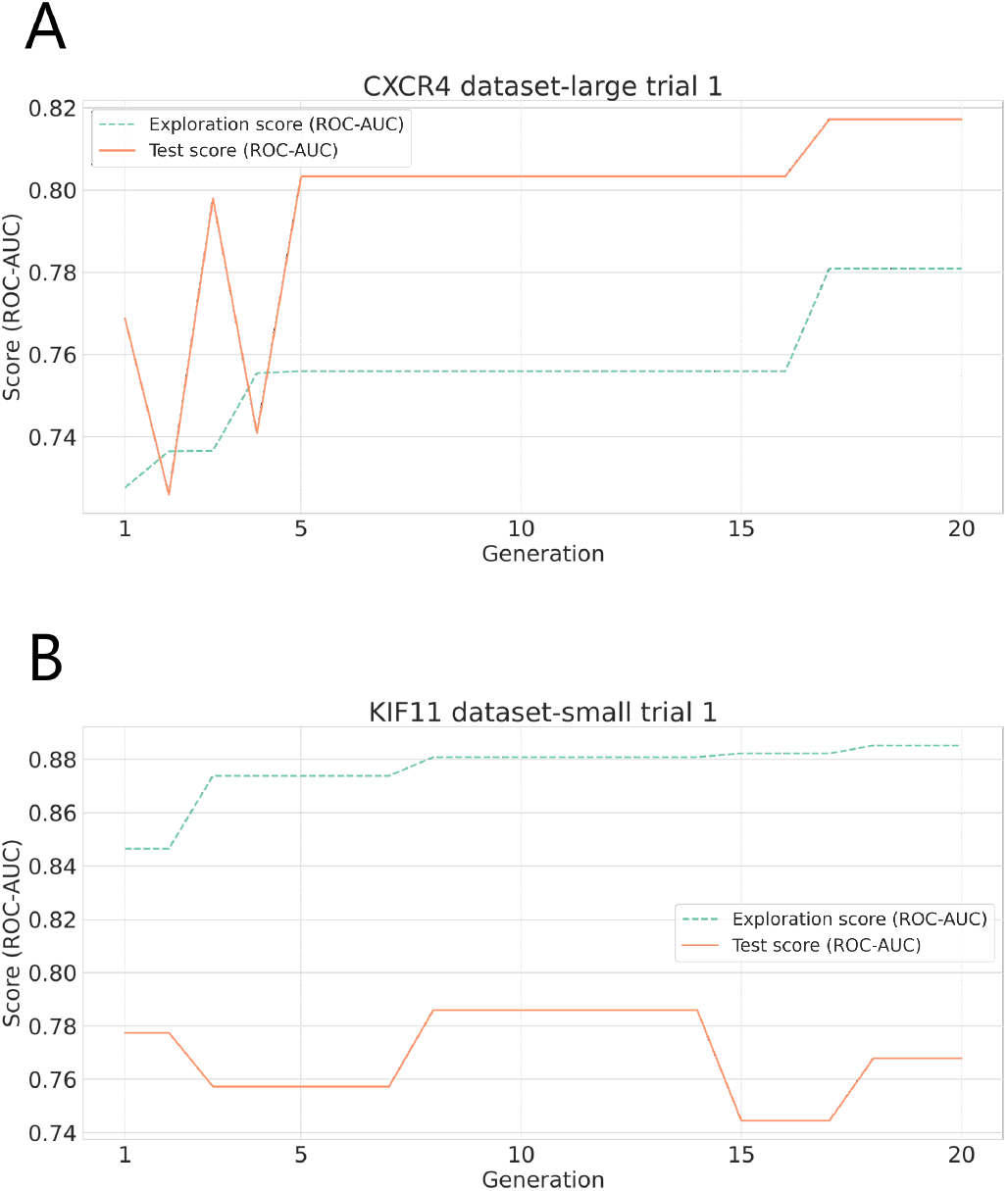
Examples of the transition of exploration in a genetic algorithm. A shows trial 1 results for CXCR4 (dataset-large). B shows trial 1 results for KIF11 (dataset-small). In both cases, exploration and evaluation were performed. The exploration score is the ROC-AUC from docking simulations using the exploration set of ligands, and the test score is the ROC-AUC from docking simulations using the test set of ligands.

On the other hand, when the number of known active compounds was small, as in dataset-small, the algorithm sometimes proceeded effectively, yet in other cases, large discrepancies arose between the exploration set and the test set (Figure 4B), causing the exploration to move in an incorrect direction or stagnate. These outcomes likely stem from the limited ligand information being inadequate to infer genuine screening performance; in such cases, the genetic algorithm might have a counterproductive effect.

### 3.3. Distribution of the explored predicted structures

We calculated the RMSD between residues within 8 °A of the ligand in the explored predicted structures and those in the apo and holo PDB structures, then visualized their distribution in a two-dimensional plot. For ABHD6 and HIPK3, no apo structure is registered in the PDB; thus, we instead used the default AlphaFold2 predicted structures. An example of the resulting distribution is shown in Figure 5.

**Figure 5:**
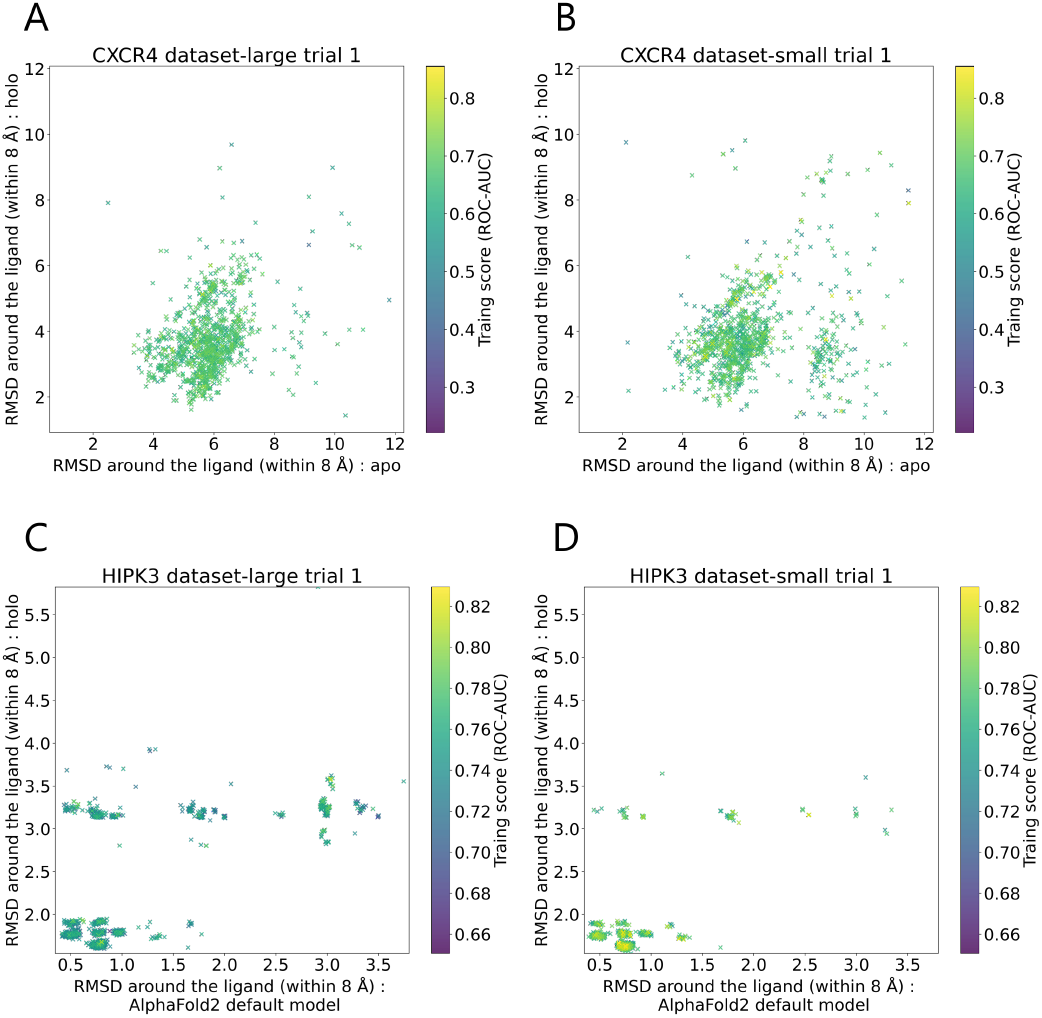
Examples of the distribution of explored AlphaFold2 predicted structures. caption A shows trial 1 results for CXCR4 (dataset-large). caption B shows trial 1 results for CXCR4 (dataset-small). caption C shows trial 1 results for HIPK3 (dataset-large). caption D shows trial 1 results for HIPK3 (dataset-small).

It is known that in a standard run of AlphaFold2, the predicted structure exhibits a single conformation [22]; however, as shown in Figure 5, the introduction of MSA mutations enables the generation of a broad conformational space in AlphaFold2 predictions. This suggests that MSA mutations are effective in enhancing the structural diversity of AlphaFold2 predicted structures. Additionally, when comparing Figure 5A and B, and C and D, the conformational space of the predicted structures did not shrink even when fewer ligands were available for exploration, indicating that the range of conformations the method can explore is not limited by the number of ligands used.

### 3.4. Transition of MSA mutations

We analyzed how the optimal MSA mutations changed in each attempt of the genetic algorithm. An example is depicted in Figure 6. From Figure 6, it can be seen that as the generations progress, mutations in specific residues gradually become fixed, indicating that the area around the previously discovered optimal MSA mutations continues to be explored. This observation implies that the genetic algorithm is functioning appropriately by refining its exploration for more suitable MSA mutations based on the best solutions identified so far.

**Figure 6:**
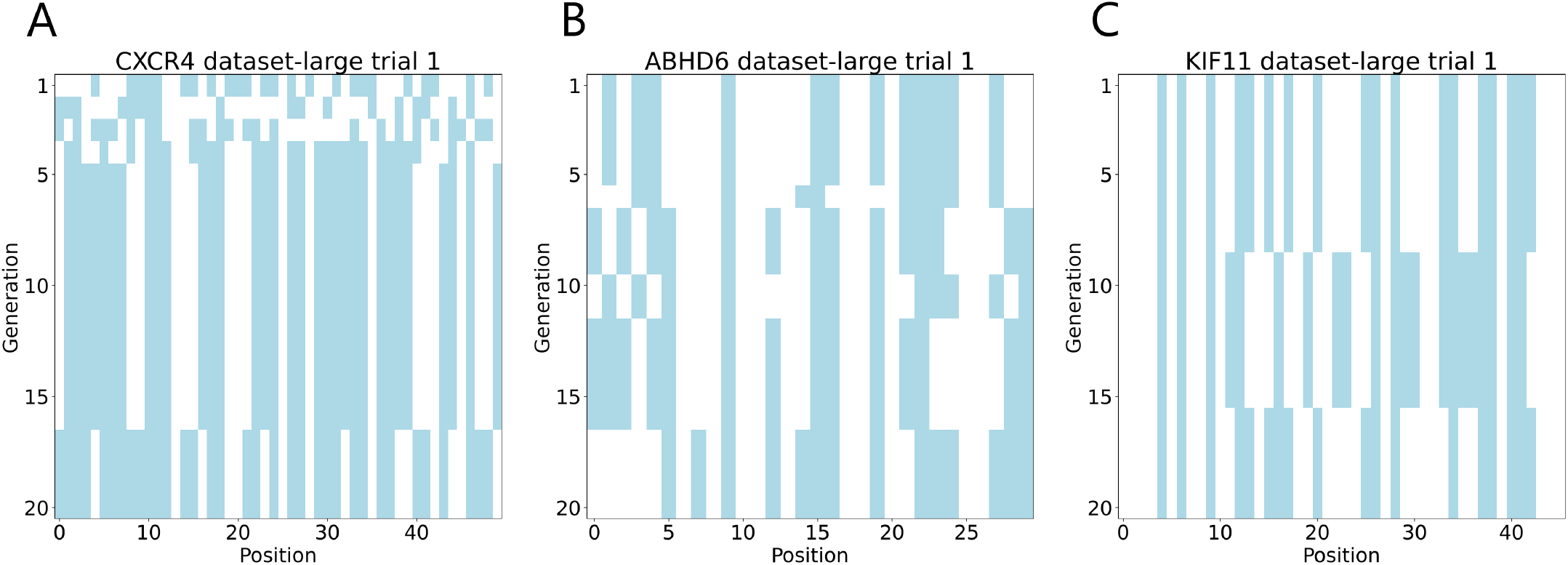
Examples of MSA mutation transitions. The vertical axis represents the generations of the genetic algorithm, while the horizontal axis shows the target residues for alanine substitutions arranged by residue number. White indicates no mutation, and blue indicates an alanine substitution.

### 3.5. Comparison of predicted structures explored by genetic algorithms and PDB Structures

We compared the tertiary structures of the best predicted structures obtained via the genetic algorithm with the holo PDB structures in order to evaluate which structural features influenced screening performance. We performed structural superposition in PyMOL, highlighting residues within 8 °A of the ligand as a binding site. The predicted structures used for comparison were those obtained in dataset-large. An example is shown in Figure 7.

**Figure 7:**
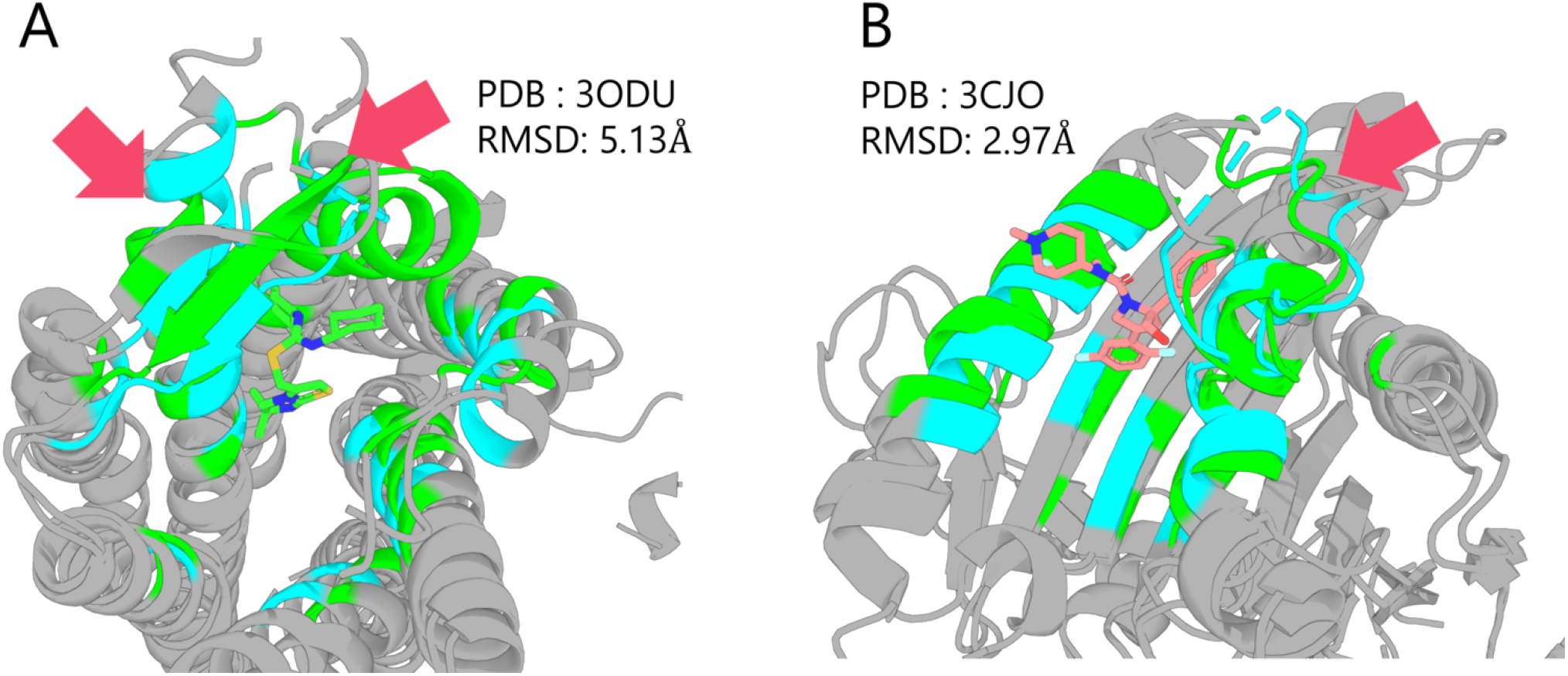
Examples of the comparison between predicted structures explored by a genetic algorithm and holo PDB structures. The predicted structure is shown in blue, and the PDB structure is shown in green. RMSD is calculated within 8 °A around the ligand.

CXCR4 (Figure 7A) exhibited a substantial improvement in ROC-AUC (0.617 to 0.793) relative to its PDB structure. A particularly large structural difference was observed in the helices surrounding the ligand, indicating that the search was able to explore configurations significantly different from those of the PDB structure.

For KIF11 (Figure 7B), no improvement was observed relative to the PDB structure (0.877 to 0.849). The main difference appeared in a loop region highlighted by arrows. AlphaFold2 is reported to struggle with accurately predicting long loops of over 20 residues [43], and the loop in question spans 17 residues. Hence, it represents a region that AlphaFold2 inherently finds challenging to predict; although the exploration identified a conformation close to the PDB structure, this difficulty seems to have led to reduced ROC-AUC performance. Nevertheless, our method improved performance over the AlphaFold2 (highest on the test set) structure (ROC-AUC: 0.834), implying that the proposed method still yields benefits for targets that are difficult for standard AlphaFold2 predictions.

### 3.6. Applicability of the proposed method to novel target

As shown in Figure 3, although ABHD6 and HIPK3 were not included among the training targets for AlphaFold2, AlphaFold2 was still able to generate predicted structures that achieved high screening performance for these proteins. This confirms the effectiveness of AlphaFold2 predicted structures in SBVS for novel targets.

Furthermore, for ABHD6 and HIPK3, the proposed method was able to explore structures that are more suitable than both the PDB structures and the average AlphaFold2 predicted structures. This highlights the applicability of the proposed method for novel targets.

### 3.7. Comparison with AlphaFold3

AlphaFold3 [44], released by DeepMind in 2024, is the successor to AlphaFold2. One of its notable features is its ability to predict protein–ligand complexes; given that holo structures tend to be more suitable for SBVS, AlphaFold3 has the potential to readily generate structures well-suited for screening. Accordingly, we evaluated the predictive performance of protein–ligand complex structures generated by AlphaFold3 under straightforward conditions and compared the results to those obtained by our proposed method.

The experimental procedure was as follows. First, we selected 10 representative ligands from the set of known active compounds by clustering. Next, for each representative compound, we ran AlphaFold3 five times and selected as the final predicted structure the one with the highest average pLDDT. We then conducted docking simulations on these predicted structures and evaluated their screening performance. The results for dataset-large are shown in Figure 8. It should be noted that AlphaFold3 was evaluated on a total of 10 structures, whereas the other methods were evaluated on 1,100 structures.

**Figure 8:**
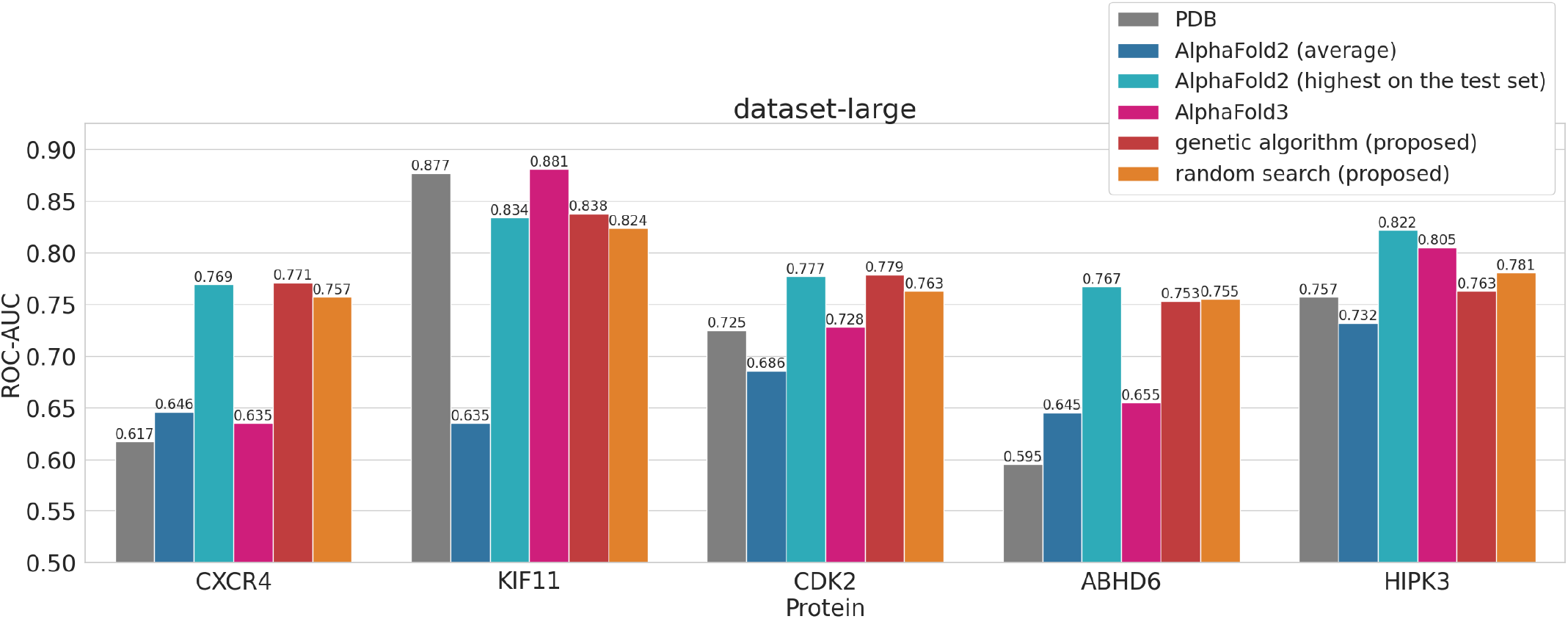
Comparison of methods based on the achieved screening performance (ROC-AUC), including AlphaFold3. The dataset-large results are shown.

From Figure 8, it is apparent that the screening performance using AlphaFold3-predicted structures varies significantly depending on the target. Specifically, for KIF11 and HIPK3, AlphaFold3 tended to yield structures out-performing other methods. Conversely, its performance in CXCR4, CDK2, and ABHD6 was markedly lower than that of the other approaches. A notable commonality is that KIF11 and HIPK3 are targets for which screening performance using PDB structures is high, whereas CXCR4, CDK2, and ABHD6 are targets for which screening performance using PDB structures is low. This observation suggests that AlphaFold3’s complex predictions may rely heavily on PDB structures. Thus, when screening performance using PDB structures is high, AlphaFold3’s predicted structures may be advantageous. Conversely, when screening performance using PDB structures is low, our proposed method, which is better at broadly exploring the conformational space, might be more suitable. Nevertheless, given that AlphaFold3’s conformational sampling for predictions remains relatively underdeveloped, additional research is required to verify the validity of these hypotheses.

## 4. Conclusion

In this study, we aimed to develop a methodology for achieving high-precision and versatile SBVS by leveraging AlphaFold2 predicted structures. Specifically, we evaluated the impact of various alanine substitution patterns in the MSA on the AlphaFold2 predicted structures using docking simulation scores with exploration compounds. Based on these results, we explored for alanine substitution patterns that can generate predicted structures suitable for SBVS—structures that enable the accurate evaluation of novel compounds. For the score-based exploration methods, both random search and a genetic algorithm were examined.

The results confirmed that, in all cases, the predicted structures explored via the proposed method were more suitable for SBVS than the average AlphaFold2 predicted structures. Furthermore, for the four targets CXCR4, CDK2, ABHD6, and HIPK3, the predicted structures explored via the proposed method tended to be more suitable for SBVS than the PDB structures. In addition, the screening performance using the explored predicted structures tended to be higher as the number of ligands available for exploration increased. In the comparison between the proposed methods, when a sufficient number of known active compounds (30 or more) was available, exploration using the genetic algorithm proved to be effective; conversely, when there were few active compounds, it appears more appropriate to employ random search. Moreover, for ABHD6 and HIPK3—which are not included among AlphaFold2’s training targets—the proposed method was able to explore structures that were more suitable for SBVS than both the PDB structures and the average AlphaFold2 predicted structures. This demonstrates the applicability of the proposed method to novel targets.

As future work, we note that both AlphaFold2 based structure prediction and protein–ligand docking simulations remain computationally intensive, indicating the need for further optimization and cost reduction for large-scale exploration. Another issue that emerged is the discrepancy between rising scores during exploration and actual performance drops. Improving the scoring function may help address this problem. Finally, AlphaFold3 holds significant promise for generating structures suited to SBVS through direct protein–ligand complex prediction, and exploring an evolutionary approach using AlphaFold3 represents an intriguing avenue for future work.

## Declaration of Generative AI and AI-assisted technologies in the writing process

No declarations were made.

## Author contributions

Keisuke Uchikawa: Conceptualization, Methodology, Software, Validation, Formal analysis, Investigation, Resources, Data curation, Visualization, Writing – original draft. Kairi Furui: Methodology, Software, Investigation, Writing – review & editing. Masahito Ohue: Conceptualization, Methodology, Formal analysis, Investigation, Writing – review & editing, Supervision, Project administration, Funding acquisition.

## Acknowledgment

Computational experiments were performed using the TSUBAME 4.0 supercomputer at the Institute of Science Tokyo.

## Footnotes

1 Note that in this study, a ‘holo’ structure is defined as a crystal structure in which any ligand is bound at the designated binding site, whereas an ‘apo’ structure is defined as a crystal structure in which no ligand is bound at that site. Under this definition of an apo structure, we do not consider ligand binding at sites other than the designated binding site. This wording differs from the generally definition of an apo structure.

