## Supporting Information for "Leveraging AlphaFold2 Structural Space Exploration for Generating Drug Target Structures in Structure-Based Virtual Screening"

### S1. AlphaFold2 Configuration

The following configuration of AlphaFold2 was used in this study:

1) **--random_seed** 1

2) **--max_seq** 16

3) **--max_extra_seq** 32

4) **--num_recycle** 1

5) **--model_order** 1

Furthermore, to ensure the reproducibility of the experimental results, the TensorFlow environment variable `TF_CUDNN_DETERMINISTIC` was set to 1 (`TF_CUDNN_DETERMINISTIC=1`), ensuring that identical inputs consistently produce identical outputs.

### S2. Genetic Algorithm Configuration

Table S1 presents the settings of the genetic algorithm used in this study. Various settings used in this study were determined to be optimal based on preliminary experiments. The rationale for each configuration parameter is presented below.

- **Population Size: 50**
  The population size was set to 50 to ensure diversity within the algorithm while keeping computational costs manageable. This allows for adequate coverage of the search space.
- **Number of Generations: 20**
  The number of generations was set to 20 to balance computational cost and convergence. This setting ensures that computation time is appropriately controlled while achieving optimal results.
- **Mutation Rate: 0.05**
  The mutation rate was set to 0.05 to maintain diversity in the search while preventing performance degradation due to excessive mutations.

*Corresponding author

Table S1: Genetic algorithm configuration

| Setting | Value |
|---|---|
| Population Size | 50 |
| Number of Generations | 20 |
| Mutation Rate | 0.05 |
| Crossover Probability | 0.9 |
| Mutation Method | bit-flip |
| Selection Method | elite preservation (2 individuals) +<br>tournament selection (tournament size = 5) |
| Crossover Method | two-point crossover |
| Scoring Method | ROC-AUC |
| Initial Population Generation Method | One individual generated without mutation and<br>149 individuals generated with a per-gene mutation probability of 0.5 |

- **Crossover Probability: 0.9**
  A high crossover probability of 0.9 was chosen to maximize the transfer of superior genes to the next generation while providing ample opportunities to generate new individuals.
- **Mutation Method: bit-flip**
  Bit-flip was adopted as the mutation method because directly switching genes between 1 and 0 allows for effective modification of genetic information.
- **Selection Method: elite preservation (2 individuals) + tournament selection**
  By preserving the top two elite individuals, this selection method guarantees that high-fitness individuals are retained in the next generation. Additionally, the use of tournament selection (with a tournament size of 5) helps maintain diversity while applying sufficient evolutionary pressure for convergence.
- **Crossover Method: two-point crossover**
  Two-point crossover was selected because it allows for the generation of diverse combinations between different parent individuals, thereby expanding the search space.
- **Initial Population Generation Method: One individual generated without mutation and 149 individuals generated with a per-gene mutation probability of 0.5**
  Given the short number of generations, the performance of the initial population significantly impacts the results. By increasing the number of initial individuals and including one individual without mutations, the method ensures diversity from the outset and enhances the stability of the search process.

### S3. Information on the Targets Used

Table S2 presents the number of known active and inactive compounds utilized for each target, along with their corresponding PDB structure IDs. The apo PDB structures were obtained using Apo–Holo Juxtaposition (AHoJ) [1].

### S4. Software Environment

The versions of the software utilized in this study are listed below.

- LocalColabFold (ColabFold) [2]: 1.5.5
- PyMOL Open-Source: 2.5.0

Table S2: Number of known active and inactive compounds and holo/apo PDB IDs for the targets

| Target | #Actives | #Inactives | Holo PDB ID | Apo PDB ID |
|---|---|---|---|---|
| CXCR4 | 122 | 3,414 | 3ODU | 4RWS |
| KIF11 | 197 | 6,912 | 3CJO | 6TA4 |
| CDK2 | 798 | 28,328 | 1H00 | 1W98 |
| ABHD6 | 111 | 5,565 | 7OTS | - |
| HIPK3 | 43 | 2,450 | 7O7J | - |

- Uni-Dock [3]: 1.1.1
- AutoDockTools: 1.5.7
- Open Babel: 3.1.1
- RDKit: 2023.09.6
